# Ascitic bacterial composition is associated with clinical outcomes in cirrhotic patients with culture-negative and nonneutrocytic ascites

**DOI:** 10.1101/322800

**Authors:** Yanfei Chen, Jing Guo, Ding Shi, Daiqiong Fang, Chunlei Chen, Lanjuan Li

## Abstract

**Background:** Ascites bacterial burden is associated with poor clinical outcomes in patients with end-stage liver disease. However, the impact of ascitic microbial composition on clinical course was still not clear. In this study, the ascitic microbiota composition of 100 cirrhotic patients with culture-negative and nonneutrocytic ascites were researched. **Results:** By characterizing the ascitic microbial composition, two distinct microbial clusters were observed, Cluster 1 (86 patients) and Cluster 2 (14 patients). Cluster 1 showed lower microbial richness than Cluster 2. At the phylum level, Cluster 1 had greater abundance of *Bacteroidetes* and *Firmicutes*, but less abundance of *Proteobacteria* and *Actinobacteria* than Cluster 2. At the family level, family *Bacteroidales S24-7* group, *Prevotellaceae, Lachnospiraceae, Lactobacillaceae, Rikenellaceae*, and *Vibrionaceae* were found over-represented in Cluster 1. And family *Acetobacteraceae, Erysipelotrichaceae, Rickettsiaceae*, and *Streptococcaceae* were found enriched in Cluster 2. The levels of plasma cytokine IL-17A, IL-7, and PDGF-BB were found significantly higher in Cluster 1 than in Cluster 2. There were four OTUs closely correlated with plasma cytokines, which were OTU 140 and OTU 271 (both from *Bacteroidales S24-7* group), OTU 68 (*Veillonellaceae*), and OTU 53 (*Helicobacteraceae*). Patients from Cluster 1 showed significant higher short-term mortality than patients from Cluster 2. **Conclusion:** Our study demonstrated that the microbial composition of culture-negative and nonneutrocytic ascites in cirrhotic patients is associated with short-term clinical outcomes. The results here offer a rational for the identification of patients with high risk, and provide references for selective use of prophylactic methods.

## Background

Bacterial translocation (BT) is increasingly recognized as a key driver in the development of complications in end-stage liver disease (1). BT can cause infections, in particular, spontaneous bacterial peritonitis (SBP). In the absence of overt infection, BT may further stimulate the immune system and contribute to haemodynamic alterations and complications. The accepted pathogenic theory of BT postulates that bacteria escape from the intestinal lumen and reach the mesenteric lymph nodes by crossing the intestinal, subsequently disseminating to the bloodstream and the ascitic fluid (AF). The presence of bacteria in AF has been investigated as a simple way for studying BT in cirrhosis.

Selective decontamination of the digestive tract (SDD) can prevent bacterial translocation and reduce severe infections and mortality in patients with end stage liver disease (2). SDD prevents secondary bacterial colonization through application of non-absorbable antimicrobial agents in the gastrointestinal tract (3). SDD is a widely evaluated but highly controversial intervention. One of the major concern about SDD is the development of bacterial resistance (4). Besides, SDD targets at both normal bacteria and potential pathogenic bacteria. The use of SDD might disturb the gut microbial balance and cause bad consequences. Hence, the identification of patients with high risk of short-term mortality and infections could provide a rational for selective use of SDD.

By culture, only a minority of ascitic bacteria can be isolated, even in the presence of overt infection. Using culture-independent techniques, Such et al. reported that bactDNA can be commonly detected in culture-negative and nonnuetrocytic AF (5). It was further confirmed by the same group that the presence of bactDNA in patients with cirrhosis during an ascitic episode is an indicator of poor prognosis, which may related to development of acute-on-chronic liver failure at short term (6). Fagan et al. found that ascites bacterial burden and immune cell profile are associated with poor clinical outcomes in the absence of overt infection (7).

With more sensitive molecular techniques such as 16S rRNA and shotgun metagenomic pyrosequencing, depth profile of microbial communities in AF have been characterized in small sample sizes recently (8, 9). Using 16S rRNA gene pyrosequencing, Rogers et al. found that differences in structure and membership of AF microbial communities correlated with severity of liver cirrhosis (8). In their research, propidium monoazide treatment were applied to ascitic samples to characterize only viable bacteria. However, the composition of translocated bacteria, both viable and non-viable, might worth studying. Circulation bacterial fragments are found with pathological significance in patients with inflammatory bowel disease (10), end stage disease (6), and chronic hemodialysis (11). Grade of soluble inflammatory response is mainly affected by plasmatic concentration of bactDNA. And, differences in inflammatory responses were observed between gram negative and gram positive bacterial fragment translocation (12).

The aim of this prospective study has been to assess the associations between AF microbial composition and clinical outcomes in cirrhotic patients with culture-negative and nonneutrocytic ascites. We applied a combination of 16S rRNA pyrosequencing and enterotype-like cluster analysis to identify AF microbial clusters. The resulting AF microbial clusters were correlated with plasma cytokine profiles and clinical outcomes. This fact may become a relevant clinical issue since it provides a reference for identification of high risk patients, who need prophylactic SDD.

## Methods

### Patients

AF were obtained from 100 consecutive cirrhotic patients undergoing clinically indicated therapeutic or diagnostic paracentesis for ascites at the First Affiliated Hospital of Zhejiang University. Cirrhosis was diagnosed by histology or by clinical, laboratory, and/or image findings. Inclusion criteria were the presence of cirrhosis and ascites fluids. Exclusion criteria were infected AF with positive culture or > 250 polymorphonuclear, upper gastrointestinal bleeding, intake of antibiotics in previous two weeks including norfloxacin as prophylaxis of SBP, two or more of criteria of systemic inflammatory response syndrome (temperature > 38°C or < 36°C, heart rate > 90 beats/minute, respiratory rate > 20 breaths/minute, blood white blood cells < 4000 or > 12000/mm^3^) (13).

AF samples were obtained when a large volume paracentesis were needed as a part of the patient’s treatment. Paracentesis were performed under aseptic conditions following the usual procedures. AF samples for routine biochemical study were obtained. Blood samples of the same day was obtained for hematological, biochemical, coagulation and cytokine profile analysis. Both blood and AF samples were kept under aseptic conditions. All the patients in study were followed up for 90 successive days. The occurrence of death and complications during the 90 days were recorded.

### DNA extraction and 16S rRNA sequencing

AF were stored at −80 °C immediately after collected. Total bacterial DNA was extracted from AF samples using QIAamp DNA Mini Kit (Qiagen, Valencia, CA) according to manufacture’s instruction. Bacterial 16S rRNA V3-V4 region was amplified using the 343F/798R primer set (343F 5′- TACGGRAGGCAGCAG −3′, 798R 5′- AGGGTATCTAATCCT −3′). PCR reaction was performed using phusion high-fidelity PCR Mastermix (Invitrogen, Carlsbad, CA, USA) with the following condition: 95 °C for 3 min (1 cycle), 95 °C for 30 s /55 °C for 30s /72 °C for 30s (35 cycles), 72 °C for 10 min. PCR product was purified using Agencourt AMPure XP beads (Beckman coulter, Brea, CA) according to manufacture’s protocol. Pyrosequencing was conducted on an Illumina Miseq 2*300 platform according to protocols.

### Pyrosequencing data bioinformatics analysis

Raw reads were filtered according to length and quality criteria. Filter-pass reads were assembled. After assembly, chimeric sequences were removed using the Usearch software based on the Uchime algorithm (14). Operational Taxonomic Unit (OTU) was picked using de novo OTU picking protocol with a 97% similarity threshold. Taxonomy assignment of OTUs was performed by comparing sequences to Greengenes. Enterotype-like clustering was performed in R with package “BiotypeR” on Jensen-Shannon distance for the OTU-level relative abundance profile (15). The optimal number of clusters was chosen based on Calinski-Harabasz (CH) values. To determine compositional features that were differentially abundant either between clusters, LEfSe was applied (16). The R package “phyloseq” was used for alpha diversity analysis (17).

### Cytokine and chemokine measurements

The plasma levels of 27 cytokines and chemokines were measured using the Bio-Plex ProTM Human Cytokine Array 27-Plex Group I on a Luminex200TM (Luminex^®^Multiplexing Instrument, Merck Millipore) following the manufacturers’ instructions. Then, we analyzed the raw data using xPONENT 3.1 software (Merck Millipore). We assumed a value of 0.1 pg/mL for statistical purposes in cases in which the concentration was undetectable.

### Statistical analysis

Two-sided student’s t-test and Mann-Whitney U-tests was used to determine whether the differences in the alpha-diversity, cytokine and chemokine levels between groups were statistically significant. We used Spearman’s rank correlation coefficient analysis to analyze the linear correlation. The Benjamini & Hochberg method was used to control the false discovery rate for multiple testing corrections. The statistical tests and plotting were done in R with package “plyr”, “ggplot2”.

## Results

### Clinical characteristics and pyrosequencing data summary

A total of 100 cirrhotic patients were included in this AF microbial profiling study. Of the patients, 56 patients were Hepatitis-B-virus related, 15 patients were alcoholic cirrhosis, 3 patients were primary biliary cirrhosis, and other 26 patients were cryptogenic cirrhosis. Of all the patients, a total of 29 patients died in 90 days, with a short term mortality of 29%. A total of 1,595,225 high-quality sequences were produced, accounting for 98.1% of valid sequences (average sequence length 419 bp).

### Two clusters identified for ascitic microbiome

The partitioning around medoids method using Jensen-Shannon distance for the OTUs-level relative abundance profile was used to investigate whether AF microbiota can be classified into clusters. The Silhouette index for two clusters was 1.86, which indicates strong support of two clusters. The two clusters were visualized by between class analyze, and showed clear separations (Figure 1a). Two clusters were confirmed with the highest CH value, as the optimal number of clusters (Figure 1b).

**Figure 1.**
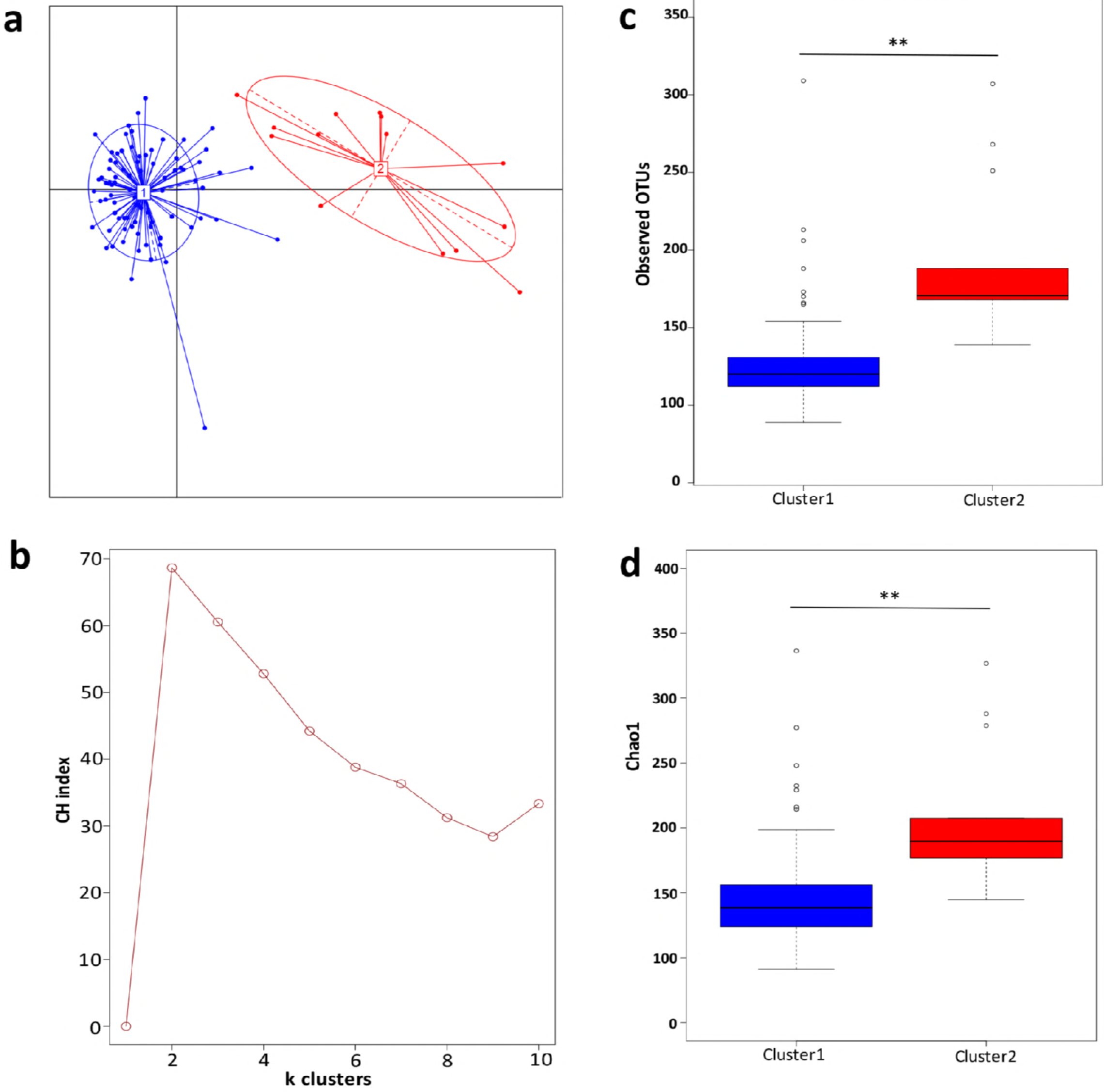
Two clusters were observed in ascitic fluids microbiota. **a**. The principal coordinate analysis of the Jensen-Shannon distance generated from the OTU-level relative abundance profiles. Samples are colored by clusters identified by the partitioning around medoids clustering algorithm. Dark blue, Cluster 1; red, Cluster 2. **b**. Two clusters were supported with the highest Calinski-Harabasz (CH) pseudo F-statistic value, as the optimal number of clusters. **c**. Boxplot comparison of the number of observed OTUs between Cluster 1 and Cluster 2. **d**. Boxplot comparison of the Chao1 index between Cluster 1 and Cluster 2. **p < 0.01 based upon Mann-Whitney U-tests with Benjamini & Hochberg correction.

When comparing the diversity index, Cluster 1 were found to have significantly lower richness than Cluster 2. After the sequencing depth normalized, the observed OTUs in Cluster 1 was significantly lower than in Cluster 2 (133±53 vs. 190±49, p=0.001) (Figure 1c), as well as the index of Chao 1 (156±69 vs. 206±53, p=0.004) (Figure 1d).

### Compositional analysis of AF microbial clusters

To identify signature taxa, we tested for significant differences among taxa displaying >1% relative abundance in the whole dataset (Figure 2). Major differences were observed between Cluster 1 and Cluster 2. At the phylum level (Figure 2a), *Bacteroidetes* and *Firmicutes* were found over-represented in Cluster 1, while *Proteobacteria* and *Actinobacteria* were found over-represented in Cluster 2.

**Figure 2.**
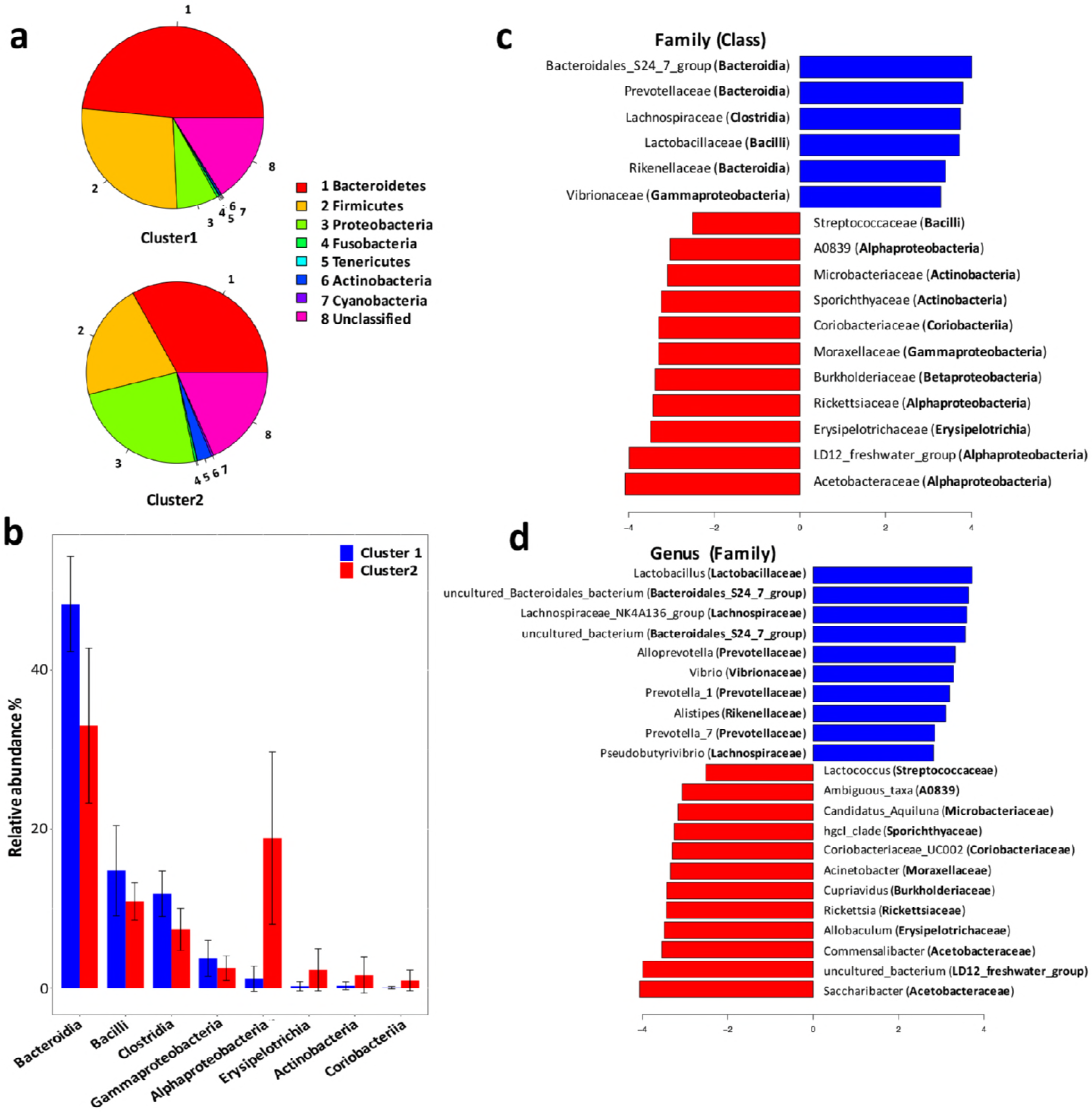
Compositional analysis of ascitic microbial clusters. **a**. Pie chart comparison of bacterial phyla represented in two clusters (upper: Cluster 1, lower: Cluster 2). **b**. Histograms showing differentially enriched bacterial classes between Cluster 1 and Cluster 2. Blue histograms, Cluster 1; red histograms, Cluster 2. **c**. LEfSe analysis revealed differentially enriched bacterial families associated either with Cluster 1 (blue) or Cluster 2 (red). **d**. LEfSe analysis revealed differentially enriched bacterial genus associated either with Cluster 1 (blue) or Cluster 2 (red).

At the class level (Figure 2b), *Bacteroidia, Bacilli, Clostridia*, and *Gammaproteobacteria* were found with significant higher relative abundance in Cluster 1 than in Cluster 2. And, *Alphaproteobacteria, Erysipelotrichia, Actinobacteria*, and *Coriobacteria* were enriched in Cluster 2.

At the family level (Figure 2c), *Bacteroidia* was linked to increased *S24-7, Prevotellaceae* and *Rikenellaceae*, while *Clostridia* expansion was linked to increased *Lachnospiraceae* in Cluster 1. Besides, *Lactobacillaceae* and *Vibrionaceae* were also found enriched in Cluster 1. The expansion of *Alphaproteobacteria* in Cluster 2 was linked to increased *Acetobacteraceae, LD12-freshwater* group, *Rickettsiaceae*, and *A0839*, while *Gammaproteobacteria* abundance was linked to increased *Moraxellaceae*.

At the genus level (Figure 2d), bacterial genus found over-represented in Cluster 1 included *Lactobacillus, Lachnospiraceae NK4A136 group, Alloprevotella, Vibrio*, and *Prevotella*. Bacterial genus *Saccharibacter, Commensalibacter, Erysipelotrichia, Allobaculum, Rickettsia*, and *Cupriavidus* were found enriched in Cluster 2.

### Plasma cytokine and chemokine levels correlated with the AF microbial compositions

We performed multiples analyses of 27 cytokine and chemokine mediators using the plasma samples collected from the cirrhotic patients. The plasma cytokine and chemokine profile was measured in 78 patients (68 patients of Cluster 1 and 10 patients in Cluster 2). The level of plasma cytokine PDGF-BB was found significantly higher in patients of Cluster 1 than in patients of Cluster 2 (12.5±18 vs. 5.2±4.2, p=0.006). Plasma cytokine IL-7 (6.39±6.31 vs. 3.96±1.87, p=0.015) and IL-17A (9.03±29.72 vs. 0.35±0.77, p=0.019) were also found significantly higher in patients of Cluster 1 than in patients of Cluster 2 (Figure 3).

**Figure 3.**
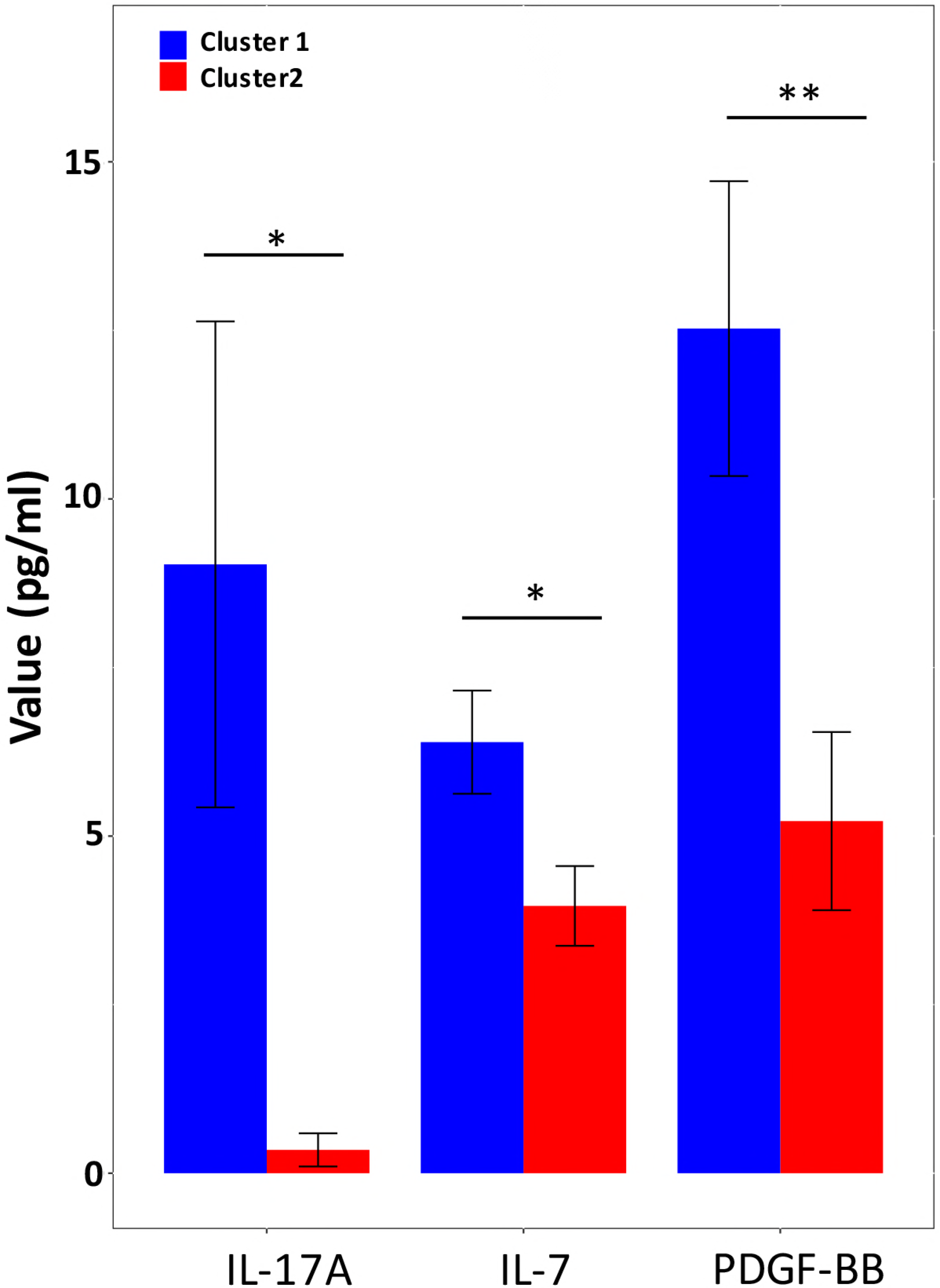
Plasma cytokine levels with statistical difference between Cluster 1 and Cluster 2. Blue histograms, Cluster 1 (n = 68); red histograms, Cluster 2 (n = 10). Significance values are indicated: * p < 0.05 **p < 0.01 based upon Mann-Whitney U-tests with Benjamini & Hochberg correction.

At the OTU level, there were 14 OTUs correlated with plasma cytokines (Figure 4). The 14 OTUs were from six bacterial families, including *S24-7 group* (6 OTUs), *Lachnospiraceae* (2 OTUs), *Rikenellaceae* (2 OTUs), *Veillonellaceae* (2 OTUs), *Helicobacteraceae* (1 OTU), and *Prevotellaceae* (1 OTU). There were four OTUs closely correlated with multiple cytokines, which were OTU 140 and OTU 271 (*S24-7 group*), OTU 68 (*Veillonellaceae*), and OTU 53 (*Helicobacteraceae*).

**Figure 4.**
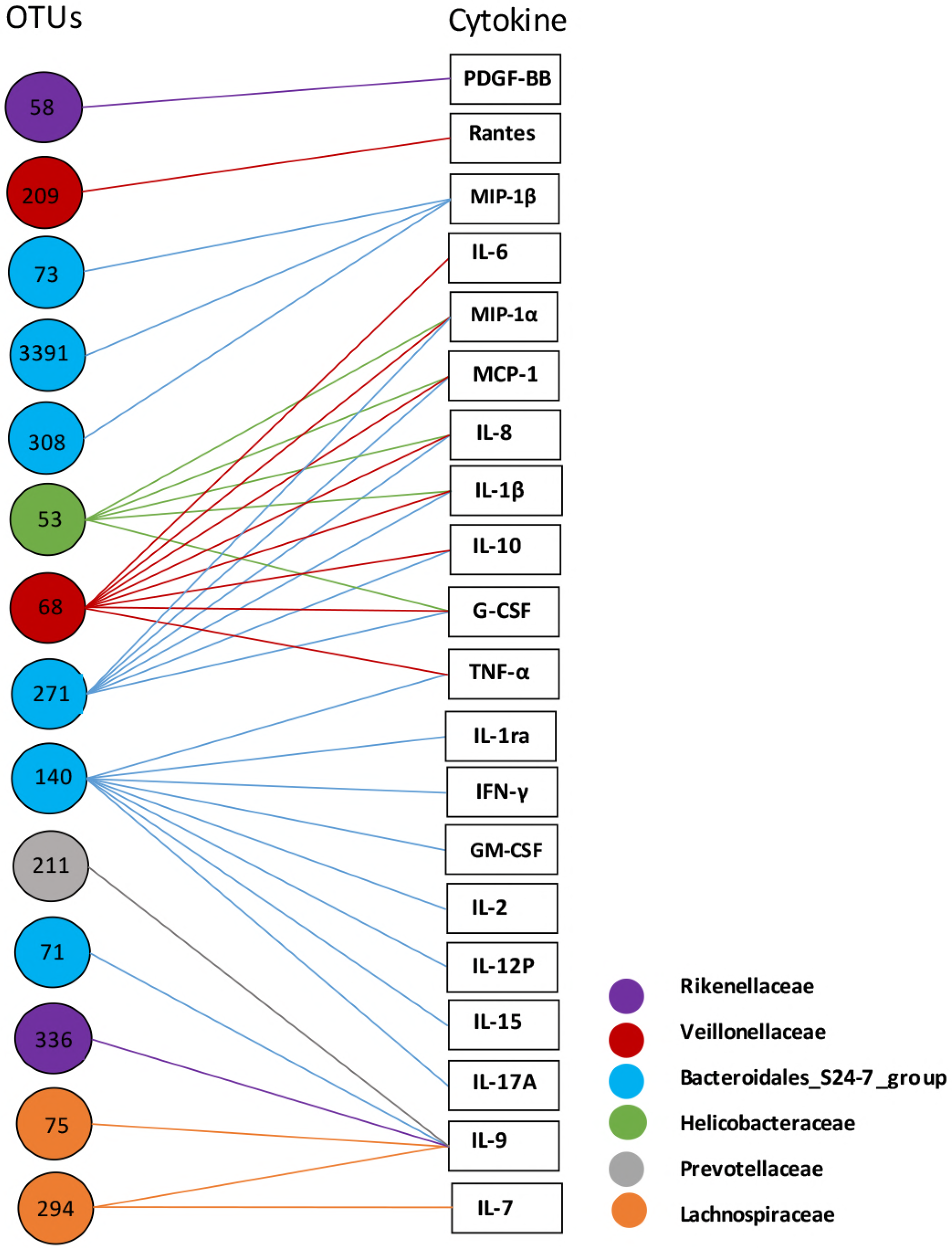
Correlation analysis of ascitic bacterial OTUs and plasma cytokines. Spearman rank correlation was performed. Only correlations with a coefficient r > 0.40 are displayed. The colors of OTU nodes and lines indicate bacterial families as labeled on the lower right.

### Cluster 1 showed higher mortality rate but lower incidence of SBP

Among the 100 patients in the study, a total of 86 patients (86%) were classified as AF Cluster 1, while other 14 patients (14%) fell into AF Cluster 2. All the patients in study were followed up for 90 successive days for occurrence of death and complications. For patients of AF Cluster 1, 29 patients died in 90 days, with a short term mortality of 33.7%. And in patients of AF Cluster 2, no patients died in 90 days. The short term mortality was significantly higher in Cluster 1 than in Cluster 2 (p=0.01) (Table 1). However, the incidence of SBP was found slightly higher in Cluster 2 than in Cluster 1 (p=0.06). Among the 86 patients in Cluster 1, only 27 patients had SBP in 90 days, with a prevalence of 31.4%. And in Cluster 2, 8 patients were diagnosed SBP in 90 days, with a prevalence of 57.1%. The severity of disease, as estimated by MELD score or Child Pugh score, was comparable between Cluster 1 and Cluster 2.

**Table1.** Clinical characteristics of patients with different ascitic microbiota clusters

| Variable | Cluster 1 (n=86) | Cluster2 (n=14) | p-value |
| --- | --- | --- | --- |
| Age (yr) | 56±11 | 54±15 | 0.7 |
| Gender (M/F) | 16/70 | 4/10 | 0.7 |
| Child-Pugh score | 9±2 | 9±2 | 0.98 |
| MELD score | 1.45±0.99 | 1.43±0.57 | 0.81 |
| 90-days mortality | 33.7% | 0% | 0.01 |
| 90-days SBP | 31.4% | 57.1% | 0.06 |

Abbreviation: SBP, Spontaneous bacterial peritonitis

## Discussion

Bacterial translocation is thought as a major mechanism of complications and mortality in end-stage liver disease. Previous studies have confirmed presence of bacterial DNA in ascites fluid, even in culture-negative and non-neutrocytic ascites. Also, positive associations between ascites bacterial burden and poor clinical outcomes was observed (7). This study demonstrated that not only the quantity of bacteria in AF but also the composition could impact the clinical outcomes of end-stage liver disease. The possible mechanism might be associated with system immune responses and cytokine expressions.

Based on the microbial composition of AF, the patients could be classified into two clusters. Cluster 1 has high mortality, and Cluster 2 has high prevalence of SBP. The group with higher mortality had more *Bacteroidetes* and *Firmicutes* than another group. *Firmicutes* and *Bacteroidetes* are the two most dominant bacterial phyla in human intestines (18). The translocation of these commensal bacteria into the abdominal cavity might indicate severe damage of intestinal mucosa barrier, which usually lead to poor outcomes. Structural and functional alterations in the intestinal mucosa that increase intestinal permeability to bacteria and its products have been described in cirrhosis (19). Altered gut microbiota profile is associated with high intestinal permeability, including less abundant of *Ruminococcaceae* and more abundant of *Lachnospiraceae* in subjects with high intestinal permeability compared with subjects with low intestinal permeability (20). Accordingly, it was observed here that ascitic microbiota of Cluster 1 have more abundance of *Lachnospiraceae* than that of Cluster 2. The findings add to the evidence that bacterial translocation from the leaky gut participate in the progress of liver cirrhosis.

Our results showed over-represented of *Alphaproteobacteria* and *Actinobacteria* in AF of Cluster 2. Both *Alphaproteobacteria* and *Actinobacteria* are common members of marine and freshwater bacterioplankton assemblages. Genomic analysis demonstrated the genome characteristics of the bacterioplankton that render them to well adapted to such nutrient and energy-limited conditions (21). One possible reason is that the intestinal permeability of patients in Cluster 2 is not severely damaged as patients in Cluster 1, and bacteria which are able to translocate to AF need greater ability to survive in nutrient-limited circumstances.

In our study, the enrichment of gut commensal bacteria in AF showed no associations with SBP. Conflict results have been demonstrated regarding the correlation between intestinal permeability and infectious complications of cirrhotics. Several previous studies found that there was no relationship of intestinal permeability with the incidence of complications (22). And, no significant difference of intestinal permeability was observed between cirrhotic patients with and without SBP (23).

Our research found that several cytokines were upregulated in patients of Cluster 1, including IL-17A, IL-7, and PDGF-BB. IL-17A is a pro-inflammatory cytokine, mainly produced by Th17 cells (24). IL-17A plays dual roles including protection host from bacterial and fungi infections, and participating in the autoimmunity diseases (25). Gut microbiota is supposed to induce Th17 differentiation and thus regulate IL-17A production and functions (26). Decreased *Firmicutes/Bacteroidetes* ratio, has been reported in systemic lupus erythematosus, that is associated with increased Th17 activation and differentiation (27). In consistent with the previous study, the AF microbiota of Cluster 1 showed decreased *Firmicutes/Bacteroidetes* ratio with increased plasma IL-17A level. The results confirmed associations between gut microbiota and Th17 differentiation. IL-17A plays important roles in fighting against pathogenic microbes using several mechanisms including induction of antimicrobial peptides, such as α-defensins, β-defensins, and lysozyme (28) (29) (30). This might explain the relative lower prevalence of SBP in patients of Cluster 1. However, Cluster 1 patients had significantly higher short-term mortality. IL-17A is able to up-regulate inducible nitric oxide synthase (iNOS), an enzyme that participates in the production of nitric oxide (31). The importance of iNOS-derived nitric oxide production in promoting BT has been evidenced experimentally in several studies. The mechanisms included that induces gastric mucosal damage, decreases the viability of rat colonic epithelial cells, directly dilates TJs in intestinal epithelial monolayers, inhibits ATP-formation and hence, increases intestinal permeability (1).

Our study demonstrated close correlations between *S24-7* family and plasma cytokines. *Bacteroidales S24-7* family is one of the substantial component of gut microbiota, which has not been successfully cultured. Multiple studies have since reported the altered abundance of *S24-7*family members in association with different physical conditions. *S24-7* is more abundant in diabetes-sensitive mice fed a high-fat diet, in particular when chow is supplemented with gluco-oligosaccharides (32). In mouse model of colitis, enrichment of *S24-7* was observed following treatment-induced remission of colitis in mice (33). Some members of the *S24-7* family are targeted by innate immune system by inducing high-affinity antigen-specific IgA responses and become highly coated with IgA (34). It was proposed that bacterial species highly coated with IgA could stimulate intestinal immunity and drive intestinal disease.

Using comparative genomics analysis, a subset of *S24-7* family was found to have urease encoding genes (35). In the intestine, bacterial urease converts host-derived urea to ammonia and carbon dioxide, contributing to hyper-ammonemia associated neurotoxicity and encephalopathy in patients with liver disease (36). Urease is also a recognized virulence factor in both bacterial and fungal infection (37). The enrichment of urease-positive species in Cluster 1, such as S 24-7 family and Lactobacillus, might suggest a possible mechanism of bacterial produced ammonia in progression of end-stage liver disease.

Bacterial class *Gammaproteobacteria* was found with significantly higher relative abundance in Cluster 2 than in Cluster 1. The *Moraxellaceae* and *Acinetabacter* were responsible for the enrichment of *Gammaproteobacteria* at the family and genus level, respectively. *Acinetobacter* species have become increasingly important nosocomial pathogens worldwide and can result in a wide range of infections, including bacteremia, pneumonia, urinary tract infection, peritonitis, among other (38). In end stage liver disease, *Acinetobacter spp*. is among the major pathogens that responsible for SBP (39). *Acinetobacter* species infections in liver transplant patients have been researched in several studies. *Acinetobacter baumannii* and *Acinetobacter Iwoffiii* were two major pathogenic species isolated (40). *Acinetobacter* infections after liver transplantation were associated with a high mortality (41). Preoperative MELD scores were more likely to be higher among the non-*Acinetobacter baumannii* compared with the *Acinetobacter* baumannii-infected group (40).

## Conclusions

Our study demonstrated that the microbial composition of AF in cirrhotic patients have correlations with short-term clinical outcomes. The possible mechanisms include intestinal permeability-dependent bacterial translocation and immune responses triggered by different translocated microbes. While the present study strengthens the association between AF microbiota and clinical outcomes, it has limitations. The intestinal microbiota and intestinal permeability of the participants was not analyzed simultaneously. We were unable to identify a direct pathway among gut dysbiosis, intestinal permeability, and immune response in end-stage liver disease. Future in vivo and in vitro studies are in need to investigate the direct correlations among intestinal permeability, translocated bacteria and immune responses.

## List of abbreviations

BT, bacterial translocation; SBP, spontaneous bacterial peritonitis; AF, ascitic fluids; SDD, selective decontamination of the digestive tract; OTU, operational taxonomic unit; CH, Calinski-Harabasz index;

## Declarations

### Ethics approval and consent to participate

The ethics committee of the First Affiliated Hospital, College of Medicine, Zhejiang University approved all the work on February 27^th^ 2014, IRB ID#2014073. All patients gave written informed consent for inclusion in the study. The data were analyzed without personal identifiers.

### Consent for publication

Not applicable

### Availability of data and material

All data generated or analysed during this study are included in this published article and its supplementary information files.

### Authors’ contributions

YFC and JG participated in the design of the study, collected biopsy samples, performed the statistical analysis, and write the paper. DS and DQF carried out the DNA extraction, and performed 16S rRNA gene PCR amplification. CLC carried out the cytokine profile analysis. LJL conceived of the study, and participated in its design and coordination and helped to draft the manuscript. All authors read and approved the final manuscript.

## Acknowledgements

Not applicable

